# Evolution of sperm morphology in a crustacean genus with fertilization inside an open brood pouch

**DOI:** 10.1101/2020.01.31.929414

**Authors:** David Duneau, Markus Möst, Dieter Ebert

**Affiliations:** Université Toulouse 3 Paul Sabatier, CNRS, IRD; UMR5174, EDB (Laboratoire Évolution & Diversité Biologique); Toulouse, France; University of Innsbruck, Department of Ecology, Technikerstrasse 25, 6020 Innsbruck, Austria; University of Basel, Department of Environmental Sciences, Zoology, Vesalgasse 1, 4051 Basel, Switzerland

**Keywords:** Sperm morphology, *Daphnia*, *Ctenodaphnia*, fertilization

## Abstract

Sperm is the most fundamental male reproductive feature. It serves the fertilization of eggs and evolves under sexual selection. Two components of sperm are of particular interest, their number and their morphology. Mode of fertilization is believed to be a key determinant of sperm length across the animal kingdom. External fertilization, unlike internal, favors small and numerous sperm, since sperm density is thinned out in the environment. Here, we study the evolution of sperm morphology in the genus *Daphnia*, where fertilization occurs in a receptacle, the brood pouch, where sperm can constantly be flushed out by a water current. Based on microscopic observations of sperm morphologies mapped on a phylogeny with 15 *Daphnia* and 2 outgroup species, we found that despite the internal fertilization mode, *Daphnia* have among the smallest sperm recorded, as would be expected with external fertilization. Despite being all relatively small compared to other arthropods, sperm length diverged at least twice, once within each of the *Daphnia* subgenera *Ctenodaphnia* and *Daphnia.* Furthermore, species in the latter subgenus also lost the ability of cell compaction by extracellular encapsulation and have very polymorphic sperm with long, and often numerous, filopodia. We discuss the different strategies that *Daphnia* evolved to achieve fertilization success in the females’ brood pouch.

## Introduction

Sexual selection is a form of natural selection acting on mating and fertilization success. Hence, sperm, the most fundamental male reproductive feature allowing egg fertilization, is expected to adapt to maximize fertilization success. In particular, two components of sperm are under selection, their number and their morphology, the latter including the associated quality. The number of sperm cells per ejaculate is known to evolve in response to several factor: risk of sperm competition for egg fertilization, cryptic female choice and, female receptacle size or the absence of receptacle (Roldan, 2019). While sperm are considered as one of the taxonomically most diverse and rapidly evolving cell types (Birkhead et al., 2009; Ramm et al., 2014), the understanding of the adaptive value of sperm morphology, such as length and shape, remains largely incomplete (Lüpold & Pitnick, 2018). Based on phylogenetic analyses across the animal kingdom, the general rule seems to be that fertilization mode (i.e. whether eggs are fertilized within or outside the female) is a key predictor of sperm length (Kahrl et al., 2021). There is a trade-off between sperm number and length (Immler et al., 2011), and therefore, the production of more sperm comes at the cost of sperm size (Parker, 1982). Sperm are generally shorter and more numerous, in species in which ejaculates are thinned out in aquatic environments, while they are longer and less numerous in species where fertilization occurs within the female (Kahrl et al., 2021; Fitzpatrick et al., 2022). This is because ejaculates are more thinned in the environment than they are within the female’s receptacle. Hence, external fertilization mode selects for higher sperm number and associated with it, smaller sperm size, to increase the chance of egg fertilization. On the contrary, internal fertilization mode reduces the thinning effect and allows for the evolution of less sperm per ejaculate and investment in higher sperm quality, e.g. longer sperm cells.

The superorder *Peracarida* is a large clade of crustaceans (e.g. *Amphipoda* and *Isopoda*) having the particularity that females have a chamber where the eggs are brooded: the brood pouch. As fertilization does occur before the females expel their eggs in the environment, such fertilization mode may be considered as internal. However, a water current generated by the filtering apparatus oxygenates the eggs in the brood pouch (Seidl et al., 2002), and it may flush out sperm. In this context, even if the fertilization is internal, we expect that the thinning effect, imposed by the water current, has an impact on the evolution of sperm length (i.e. reduces their size). Furthermore, males are likely to evolve persistence traits that allow them to avoid being flushed away. Out of the 4705 species from which we have a sperm morphology description in the SpermTree database, only 9 species belong to the superorder of the *Peracarida* (Fitzpatrick et al., 2022). Three of them have sperm described as aflagellate and the others are described as non-standard, as they have tail-like morphology which are not a flagellum. Despite being a very recent and rich database, little is known about sperm in this species rich clade with a particular fertilization mode that may affect the evolution of sperm morphology.

The water fleas or *Daphnia* are not part of the *Peracarida*, they are *Cladocera*, but they possess a brood chamber, where fertilization of sexual eggs takes place. They reproduce mostly by cyclical parthenogenesis, with sexual reproduction and thus egg fertilization being sporadic, but essential for diapause in freezing and drying habitats and for dispersal. Usually triggered by a change in environmental conditions and following periods of clonal reproduction during which females only produce genetically identical daughters, some females produce sexual (haploid) eggs while others produce males (Deng & Lynch, 1996; DeMeester & Vanoverbeke, 1999). During mating, one or two males attach to the female to fertilize eggs, which will be released by the female in her brood pouch after the male(s) departed. The brood pouch is a receptacle formed by the carapace on the dorsal side of all *Daphnia* species, where early development of clonal and sexual eggs takes place. For the latter, the cuticular structure of the brood pouch changes to form a protective case, which will be released upon molting. These resting stages contribute to genetically diverse egg-banks from which future populations can be established. There is evidence for males competing for fertilization in *Daphnia magna* (Duneau et al., bioRxiv 2020), but the extend of sperm competition in this receptacle is unknown. It is not entirely clear where fertilization exactly occurs, whether in the brood pouch or inside the oviduct exiting in the brood pouch. On a one hand, sperm are not flagellated (Wingstrand, 1978; Wuerz et al., 2017), and it is difficult to see how sperm could enter the oviduct as it is closed and eggs have to push their way out (Lee et al., 2019). On the other hand, *Daphnia* males have more or less exaggerated genital papilla, the organ used to deposit sperm in the brood chamber. When exaggerated, they could help to bring sperm close to the oviduct, which was used to argue that fertilization occurs in the oviduct (Baldass, 1941) or at its entrance, and reduces the thinning effect of the water current in the brood pouch.

Although pioneer studies have given key general descriptions to identify the main sperm structures (Delavault & Berard, 1974; Wingstrand, 1978; Zaffagnini, 1987; Wuerz et al., 2017), only little is known about morphology of aflagellated sperm in *Daphnia.* All *Anomopoda*, an infraorder including *Daphnia*, have a vacuolar type of spermatogenesis (Wingstrand, 1978), i.e. in the testes, the spermatids are enclosed in “private” vacuoles in the nutritive cells and are exocytosed into the testicular lumen after they have decreased strongly in size and matured. When compacted during maturation, they are generally small, about a few microns in length and width. Sperm of *D. magna* has been more thoroughly studied with recent technology. This representative of the subgenus *Ctenodaphnia* has longer sperm (~10 μm) encapsulated by an acellular capsule, probably to pack more sperm in the testes, and very short filopodia extending from the cell membrane (Wuerz et al., 2017). The roles of this capsule and of the filopodia are unclear (Wuerz et al., 2017). Here, we reconstructed a robust phylogeny of the *Daphnidae* using COI, 12S and 16S rRNA genes based on (Adamowicz et al., 2009; Cornetti et al., 2019) and assessed 15 species representing major clades within *Daphnia* to better describe the variation in sperm morphology in this genus and to gain insight into its evolution.

## Methods

Male *Daphnia* were either sampled from female mass cultures in the laboratory in which male production was naturally occurring as a consequence of high population density, or from females exposed to the hormone methyl farnesoate (MF, 400nM final concentration) which is known to induce male production (Olmstead and Leblanc 2002). We assessed samples covering major clades in the genus *Daphnia.* For the subgenus *Ctenodaphnia* we included *D. similis* Claus, 1876, *D. sinensis* Gu, Xu, Li, Dumont et Han, 2013, *D. lumholtzi* Sars, 1885, *D. carinata* King, 1853, *D. magna* Straus, 1820, *D. hispanica* Glagolev and Alonso, 1990, *D. dolichocephala* Sars, 1895, and *D. barbata* Weltner, 1897. For the subgenus *Daphnia*, we sampled *D. pulex* Leydig, 1860 and the European clade of *D. pulicaria* Forbes, 1893 as representatives of the *D. pulex* group sensu lato. From the same subgenus we also included samples from the *D. longispina* group sensu lato, sometimes also referred to as ‘*Hyalodaphnia*’ (Petrusek et al., 2008). This group was represented by members of the *D. longispina* complex (Petrusek et al., 2008) - namely *D. longispina* O.F. Müller, 1785,*D. mendotae* Birge, 1918, *D. galeata* Sars, 1864, and *D. dentifera* Forbes, 1893 – as well as *D. curvirostris* Eylmann, 1887. For *D. longispina* our sampling also covered the different morphotypes *“zschokkei”, “hyalina”* and *“longispina”* (Petrusek et al., 2008).

We induced male production for the *D. longispina* morphotypes *“hyalina”* and *“zschokkei”, D. dentifera, D. mendotea, D. galeata*, and *D. curvirostris* and collected naturally produced males for *D. similis, D. sinensis*, *D. lumholtzi*, *D. carinata*, *D. magna*, *D. hispanica*, *D. dolichocephala*, *D. barbata*, *D. longispina (“longispina”* morphotype), *D. pulex* and *D. pulicaria.*

To collect sperm, we exposed mature males to a 1 % nicotine solution ((-)-Nicotin 162.23 g/mol, supplier: Carl Roth, Germany) to induce ejaculation as in Duneau et al. (2012). As only mature spermatozoa are present in the testicular lumen (Wingstrand, 1978:11; Zaffagnini, 1987:277), this method allowed us to describe and measure mature sperm and avoid immature ones. Presence of filopodia on the sperm was recorded, but we did not measure filopodia length. Measurements of the greatest length of the freshly released sperm were performed with ImageJ (Open source, v. 1.5i) using photographs taken under phase contrast light at a magnification of 40x. In species with very small sperm (*D. pulex, D. pulicaria, D. dolichocephala* and *D. barbata*) we paid particularly attention that the sperm were photographed just after release from the spermiduct to reduce the possibility of degradation or to confuse them with other particles. However, it was challenging to take good photographs of them, and the measurement may be less accurate than for the other species. For example, *D. pulex* sperm is only around 2 μm in length (Xu et al., 2015). All sperm were also observed at the moment of release from the ejaculatory opening to verify that their shape corresponds to what was observed later when they settled and were photographed. We also observed sperm morphology in sea water to test that osmolarity was not affecting our results. We did not see any effect and do not report further on these observations. Drawings of male genital papilla were redrawn from published identification keys (Benzie, 2005; Popova et al., 2016).

For an ancestral trait reconstruction, we reconstructed a phylogeny based on the mitochondrial COI and 12S and, where available, 16S rRNA genes published in Adamowicz et al. (2009) and Cornetti et al. (2019) (for sample information see Supplementary Table 1). Sequences were aligned with MUSCLE (Edgar, 2004) and a maximum-likelihood tree partially constrained for the topology of the full mitochondrial genome phylogeny from Cornetti et al. (2019) was constructed with iqtree2 (Nguyen et al., 2015; Minh et al., 2020). First, the best model and best-fit partitioning scheme were inferred with ModelFinder (Chernomor et al., 2016; Kalyaanamoorthy et al., 2017) (-m MFP+MERGE; for full commands see Supplementary Text 1). We then used the best model and scheme to calculate a constrained and an unconstrained tree and compared the two trees using tree topology test with the RELL approximation using 10,000 replicates (Kishino et al., 1990), i.e. the bootstrap proportion, Kishino-Hasegawa test (Kishino & Hasegawa, 1989), Shimodaira-Hasegawa test (Shimodaira & Hasegawa, 1999), and expected likelihood weights tests (Strimmer & Rambaut, 2002)), and the approximately unbiased test (Shimodaira, 2002) (-zb 10000 -au). Since there was no significant difference between constrained and unconstrained trees, we continued with the constrained tree and calculated SH-aLRT (Guindon et al., 2010) and UFBoot (Hoang et al., 2018) support values with 10,000 replicates (-bb 10000 -alrt 10000). The constrained tree was then also used for ancestral state reconstruction using the packages phytools 0.7-70 (Revell, 2012), ape 5.4-1 (Paradis & Schliep, 2019), and geiger (Harmon et al., 2008; Pennell et al., 2014) in R version 4.0.3 (2020-10-10) (R Core Team, 2020). We obtained an ultrametric dated tree using our ML tree and multiple node calibration using the mtDNA - fossil calibration - Late Jurassic age bounds from Cornetti et al. (2019) and a strict clock model in chronos(). We then calculated mean sperm length values for each measured *Daphnia* species and added mean values for *Ceriodaphnia reticulata* and *Simocephalus* sp. from (Wingstrand, 1978). For ancestral state reconstruction, we used the sperm length data and the dated tree and fitted a Brownian motion model of evolutionary change with phytools (code, alignments and input files for tree reconstruction and ancestral state analysis are deposited on GitHub - https://github.com/markusmoest/daphnia_sp) and on Zenodo (https://doi.org/10.5281/zenodo.6992749).

## Results

Among the species we studied, the genital papilla was only exaggerated in *Daphnia magna*, but inconspicuous in the other species. Hence, we could not correlate this trait with sperm morphology. Using microscopic observations of sperm morphologies allowing to measure the maximum length of rod-shape aflagellated sperm cells, we found that sperm length varied greatly among *Daphnia* species, ranging from about 2 μm to at least 20 μm (Figure 1 and supplementary figures 1 and 2). Based on recent *Daphnia* phylogenies (Adamowicz et al., 2009; Cornetti et al., 2019), we found that there was a clear phylogenetic signal in sperm length across *Daphnia*, but length clusters are polyphyletic. Sperm length diverged at least twice in the genus *Daphnia* (Figure 1), once in the subgenus *Daphnia* and once in the subgenus *Ctenodaphnia.* We also found that the assessed species in the *D. longispina* clade, i.e. *D. longispina, D. dentifera, D. mendotae* and *D. galeata*, have entirely lost the capsule compacting sperm and had very long filopodia (Figure 2 C and D, Supplementary figures 1, 2 and 3).

**Figure 1:**
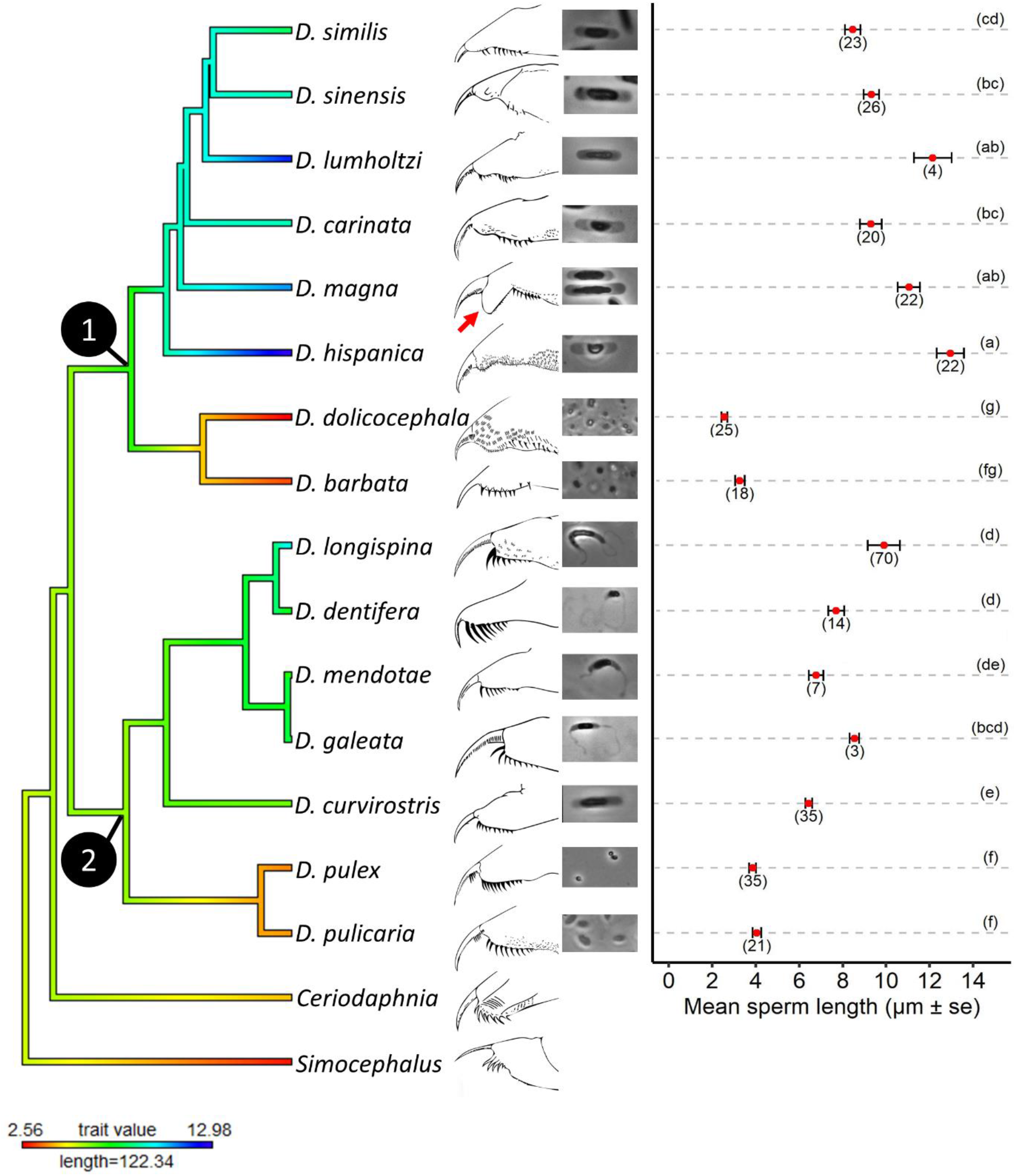
Evolution of sperm length and genital papilla morphology in *Daphnia*. Phylogenetic tree is modified from Cornetti et al. (2019) and Adamowicz et al. (2009). Color gradient represents sperm length and length of scale bar indicates to branch length in million years. Drawings represent the genital papilla of the males (redrawn from Benzie (2005) and Popova et al. (2016)). The red arrow indicates the atypical exaggerated genital papilla of *D. magna.* Photographs show an example of sperm for each species. The graph on the right represents the difference in sperm length among males. The mean sperm length was calculated from 2 to 3 individuals per species. Numbers under the mean represents the number of measured sperm. Letters on the right illustrates whether median sperm lengths are significantly different (different letters) or not (same letter) among species based on pairwise Kruskal-Wallis comparisons corrected for multiple testing using the FDR method. Numbers in phylogeny represent the node of the two main *Daphnia* clades: 1: subgenus *Ctenodaphnia*; 2: subgenus *Daphnia*.

**Figure 2:**
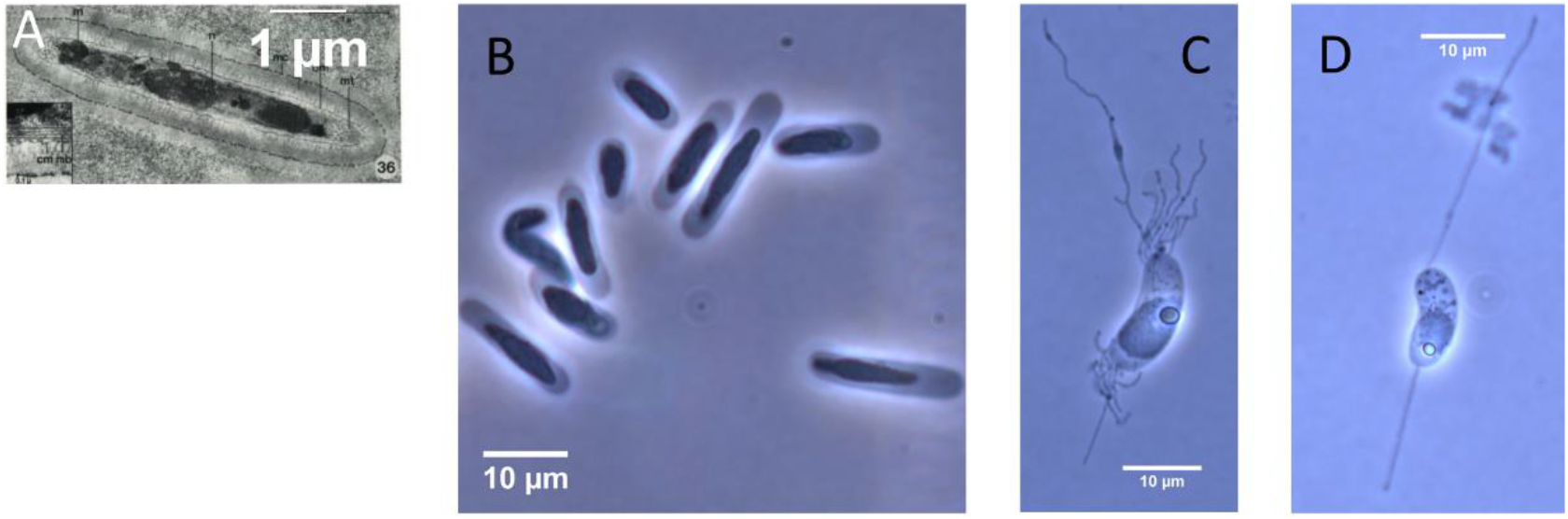
Examples of sperm morphologies in the *Daphniidae*. A- *Ceriodaphnia laticauda*, small and encapsulated elongated sperm (blue color in figure 1) representative of one of the species used as outgroup. Photo from Wingstrand (1978). B- *D. magna*, large and encapsulated elongated sperm (blue color in figure 1); C and D - *D. longispina*, two examples of typical non-encapsulated and elongated sperm with filopodia (green color in figure 1).

*Ceriodaphnia* and *Simocephalus*, both members of the family *Daphniidae* and used here as outgroups, have sperm of the vacuolar spermatogenesis type, like *Daphnia* species and all other clades in the infraorder *Anomopoda.* Their sperm have been described, based on electron microscopy, as small (2 to 6 μm), more or less rod-shaped and strongly compacted in their capsule (e.g. Figure 2A and Wingstrand, 1978:25-26). This information based on several species allowed to assume, although with caution, that the most parsimonious ancestral sperm type in the genus *Daphnia* was rather short. The subgenus *Ctenodaphnia*, except for *D. dolichocephala* and *D. barbata* who had compacted and small sperm, evolved non-compacted and elongated sperm, several times larger than the putative ancestral morphology (Figure 1 and 2B). A similar adaptation occurred in *Daphnia* s. l.. In accordance with reports from Xu et al. (2015) on *Daphnia pulex* sperm length, we observed that members of the *D. pulex* group sensu lato conserved the small and compacted sperm morphology, whereas members of the *D. longispina* group sensu lato evolved larger elongated sperm (Figure 1). Furthermore, in the same ejaculate from a single *D. longispina* male, sperm can have a strong polymorphism in the number and length of filopodia per sperm cell. The cell can have, on each side, one long filopodium or many shorter ones (Figure 2C and 2D, and Supplementary figure 3). These filopodia can be several times the length of the sperm (not measured here) (Figure 2 and Supplementary figure 1 and 3).

## Discussion

Since *Daphnia* males ejaculate in the female brood pouch, the fertilization mode is here considered to be internal. However, as a water current generated by the filtering apparatus constantly circulates through the brood pouch (Seidl et al., 2002), male *Daphnia* face the challenge to have their sperm flushed out. This phenomenon could be seen as a form of cryptic female choice, as it increases the thinning effect. When two males copulate at the same time with a female, the brood chamber is a place for direct sperm competition to occur. Interspecies variations in such sexual selection may be present and shape sperm evolution. Males of different species may have different features to increase their chance to successfully sire offspring. We speculate that when species have a brood pouch, males may be selected to: 1- produce more, but smaller sperm, 2- deposit their sperm the closest to the oviduct, 3- develop structures to reduce the chance that sperm is flushed out.

Our assessment of sperm morphology over 15 *Daphnia* species uncovered clearly structured phylogenetic variation in sperm length. It diverged at least twice, once in the subgenus *Daphnia* and once in the subgenus *Ctenodaphnia.* Our study reports that *Daphnia* have small rod-shape aflagellated sperm with the species averages ranging from 2.6 to 13 μm. Considering that sperm in arthropods measure on average 1034.4 μm (±81.7 SE and ±3051.5 SD) (based on 1394 arthropod species in SpermTree database, (Fitzpatrick et al., 2022)), our work allows to conclude that sperm are relatively small in *Daphnia.* This supports the hypothesis that the particular internal fertilization of species with brood pouch do not necessarily favor the evolution of larger sperm, most probably due to the thinning of sperm density imposed by the constant water flow.

Variation in aflagellated sperm length within each *Daphnia* subgenus was ranging from 2.6 to 13 μm in the subgenus *Ctenodaphnia* and from 3.8 to 9.9 μm in the subgenus *Daphnia.* From the 185 species with aflagellated sperm of which the length is available in the SpermTree database, the median of the averaged total length is 5 μm, with a range from 1 to 500 μm (Calculated from SpermTree database, Fitzpatrick et al., 2022). Two *Daphnia* species from each subgenus had mean sperm length below this median length of 5 μm, while all other species were above this. Part of the size of sperm is due to an extracellular compaction by an extracellular vacuole which takes place before the mature sperm is released into the spermiduct (Wingstrand, 1978). As most sperm production happens when males are juveniles, the total amount of sperm material is limited by the size of the spermiduct (Wuerz et al., 2017). By limiting the total amount of sperm material stored, this constraint may put selection on the degree of compaction allowing to store more cells in the duct. In such a case, *Daphnia* species with “large” sperm may had relatively relaxed selection on sperm compaction, in comparison to species with “small” and very compacted sperm. We propose that although, fertilization inside a brood pouch favors smaller sperm probably to favor numbers, the evolutionary changes in sperm length we observed may in part be underlined by a change in the mechanism of cell compaction before maturation due to relaxed selection for extreme compaction in some species.

Another way to increase the chance of fertilization in the brood pouch is to deposit the sperm closer to the oviduct. Few *Daphnia* species are known to have exaggerated genital papilla (Flössner, 2000; Benzie, 2005). We wondered if the penis-like apparatus, which could help to position the sperm closer to the oviduct and thus reduce the thinning effect, could correlate with sperm size. However, we found that despite the phylogenetic signal in sperm length, only one species included in our analysis, *D. magna*, has an exaggerated genital papilla. Hence, genital papilla morphology cannot be used to explain the phylogenetic structure we observed. However, strength of sexual selection is a function of how often male ejaculates compete for fertilization, in particular via direct sperm competition and sperm competition through cryptic female choice. If exaggerated genital papilla evolved under such male-male competition and reduce the thinning effect by improving the chance that sperm are deposited close to the oviduct, one can speculate that this trait evolves when strong sexual selection occurs in the system. It is difficult to estimate the intensity and frequency of sexual selection in the system, especially for each species. The induction and frequency of sexual reproduction in *Daphnia* depends on the species and the environment and thus constitutes a locally adapted trait (Roulin et al., 2013). In unstable and short-lived habitats, such as small rockpools or ponds in strongly seasonal environments like deserts and arctic sites, few asexual generations occur before diapause recommences. In stable environments, such as large lakes and ponds in temperate mild climatic regions, many asexual generations may occur before the next sexual generation and most individuals will never go through a sexual cycle. Traditionally, the latter case received more attention by *Daphnia* researchers, leading to the biased impression that sexual reproduction, and thus the occurrence of males, is generally rare. However, in *Daphnia magna*, there is a large heterogeneity among populations. In unstable environments, males can be periodically abundant, and several males can be found copulating at the same time with a female, suggesting that sexual selection can be strong, maybe more than in most other species (Duneau et al., bioRxiv 2020). Noticeably, *D. magna* is the only species here that have exaggerated genital papilla and the only species for which matings with multiple males have been reported (Duneau et al., 2020). One could wonder if the evolution of exaggerated genital papilla could have evolved in *D. magna* due to selection to improve sperm deposition close to the oviduct and reduce the thinning effect in a species under strong sexual selection. More work is needed to strengthen this hypothesis. One way would be to compare the variation in genital papilla size between stable and unstable *D. magna* populations, as those populations should differ in the intensity of sexual selection. We would expect that individuals from stable populations would have less exaggerated genital papilla than those from unstable populations.

In an open brood pouch, sperm would gain an advantage if they could attach to the inner lining of the pouch, so the water current will not flush them out. We did not observe a structure supporting the idea that encapsulated sperm can attach to the inner lining. However, it is possible that, like in crayfish (Niksirat et al., 2014), the extracellular capsule might break post-copulation and reveal filopodia which could attach to surfaces. In *Daphnia magna*, such filopodia have been reported inside the extracellular vacuole (Wuerz et al., 2017), but they are very small and difficult to resolve, even with electron microscopy. Our phylogenetic analysis of sperm morphology revealed a monophyletic clade (i.e. *D. longispina* species complex) which evolved an apparent loss of the capsule and the gain of long, sometimes numerous, filopodia of diverse shapes. The lengths of these filopodia may be a multiple of the length of the sperm cell (Fig. 2). We did not observe any movement of the filopodia and therefore consider it is unlikely that the flagella are used to move towards the eggs. The filopodia are very flexible and can be forked. But, as sperm features are expected to be adaptations to their specific fertilization environment (Pitnick et al., 2009), it is reasonable to assume that the filopodia play a role in fertilization. This role may be related to the fusion with the oocyte, but it may also reduce the chances to become flushed out from the brood pouch. Hence, we suggest that the loss of the capsule may have evolved from the reduced selection for compaction and the advantage to improve fusion with the oocyte and attach to the brood pouch.

## Acknowledgements

We thank Jürgen Hottinger for help in the laboratory, Luca Cornetti for help with the alignment files from his study on *Daphnia* phylogeny, and Jeremias Brand, Vitor Faria, Scott Pitnick, Lukas Schärer and Axel Wiberg for discussions about the study. We thank Birgit Schlick-Steiner and the Molecular Ecology group for hosting DD during his research stay. Version 4 of this preprint has been peer-reviewed and recommended by Peer Community In Evolutionary Biology (https://doi.org/10.24072/pci.evolbiol.100145)

## Data, scripts and codes availability

Data are available online: https://doi.org/10.5281/zenodo.6991181

Script are available online: https://doi.org/10.5281/zenodo.6992749

## Appendix

**Supplementary figure 1:**
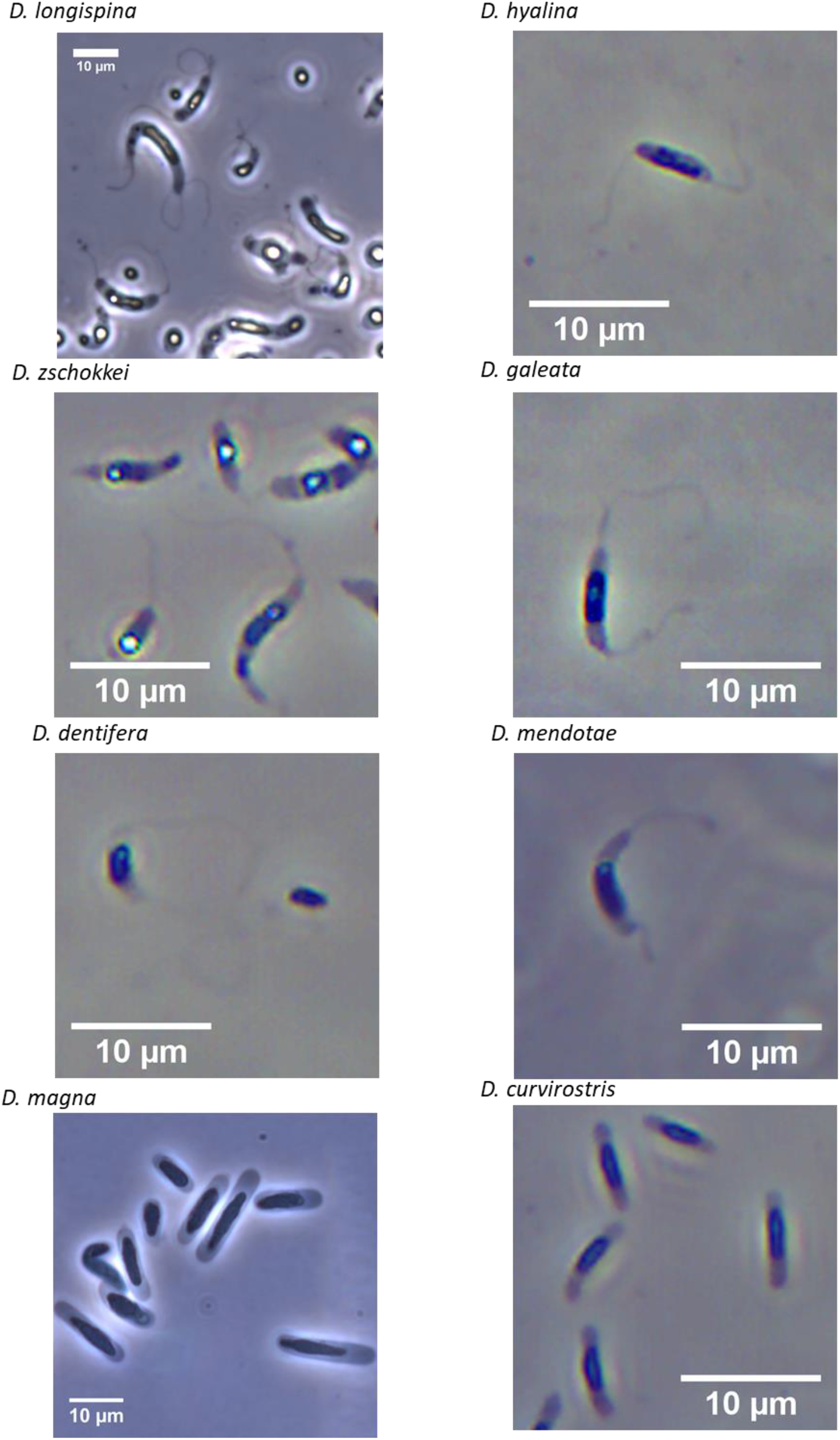
Illustration of *Daphnia* sperm morphology by species.

**Supplementary figure 2:**
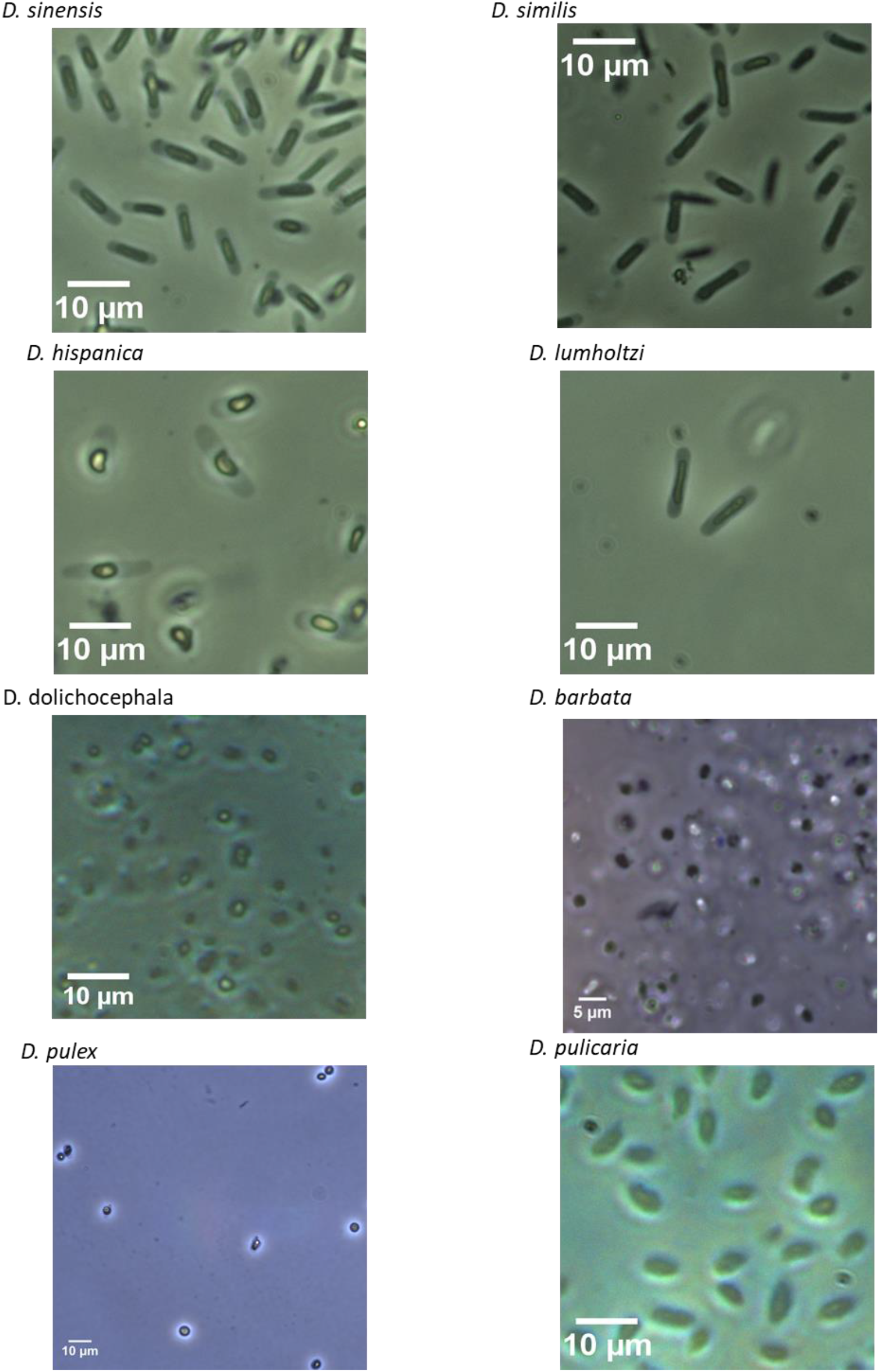
Illustration of *Daphnia* sperm morphology by species.

**Supplementary figure 3:**
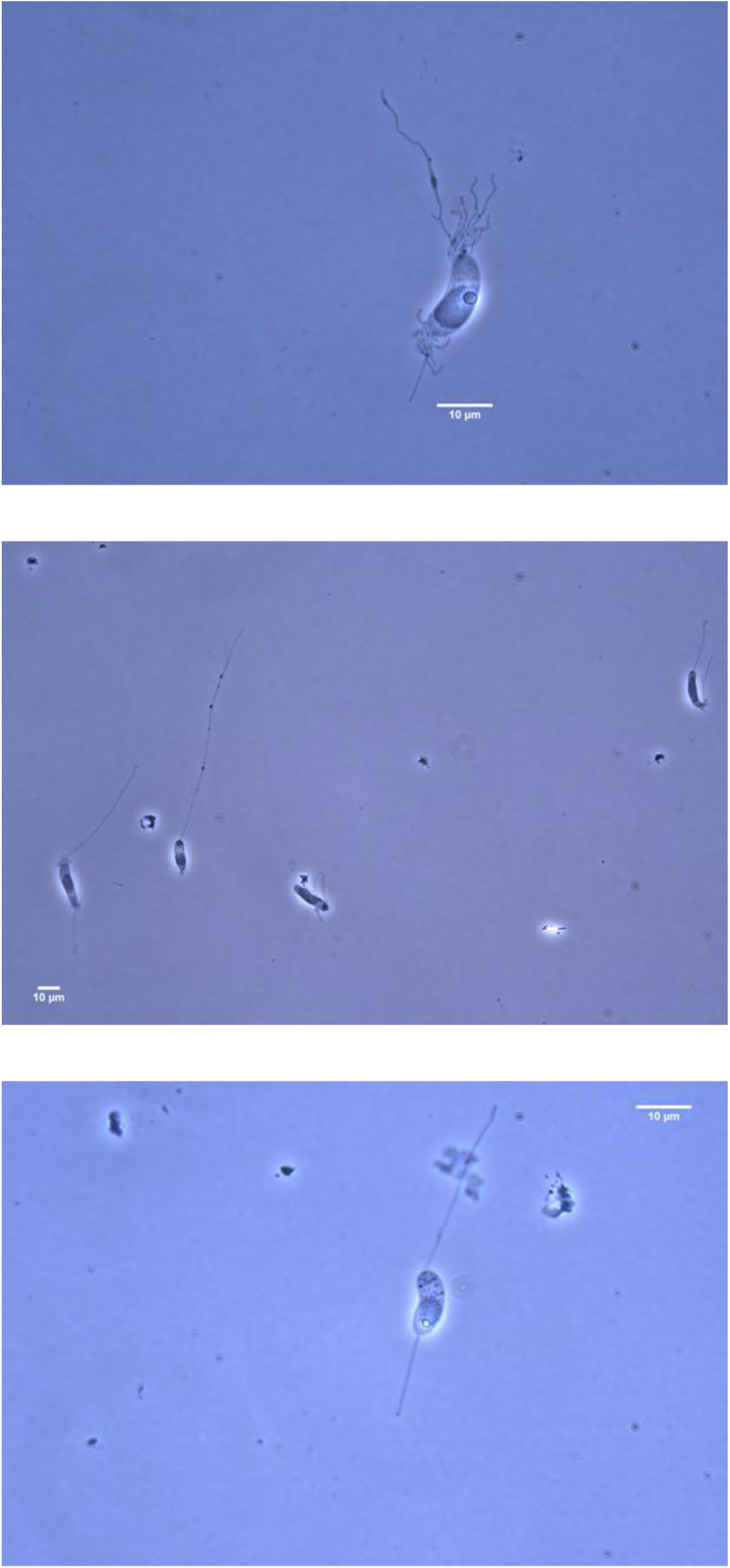
Examples of *Daphnia longispina* sperm morphologies.

## References

Adamowicz, S.J., Petrusek, A., Colbourne, J.K., Hebert, P.D.N. & Witt, J.D.S. 2009. The scale of divergence: A phylogenetic appraisal of intercontinental allopatric speciation in a passively dispersed freshwater zooplankton genus. Molecular Phylogenetics and Evolution 50: 423–436. https://doi.org/10.1016/j.ympev.2008.11.026

Baldass, F. 1941. Entwicklung von Daphnia pulex. In: Abteilung Fur Anatomie und Ontogenie der Tiere.

Benzie, J.A.H. 2005. Cladocera: The genus Daphnia (including Daphniopsis). Backhuys Publishers, Leiden, The Netherlands.

Birkhead, T.R., Hosken, D.J. & Pitnick, S. 2009. Sperm biology: an evolutionary perspective, Academic P. Elsevier.

Chernomor, O., Von Haeseler, A. & Minh, B.Q. 2016. Terrace aware data structure for phylogenomic inference from supermatrices. Systematic Biology 65: 997–1008. https://doi.org/10.1093/sysbio/syw037

Cornetti, L., Fields, P.D., Van Damme, K. & Ebert, D. 2019. A fossil-calibrated phylogenomic analysis of Daphnia and the Daphniidae. Molecular Phylogenetics and Evolution 137: 250–262. https://doi.org/10.1016/j.ympev.2019.05.018

Delavault, R. & Berard, J.J. 1974. Ultrastructural study of spermatogenesis in Daphnia magna Straus (Entomostraca, Branchiopoda, Cladocerae).

DeMeester, L. & Vanoverbeke, J. 1999. An uncoupling of male and sexual egg production leads to reduced inbreeding in the cyclical parthenogen Daphnia. Proceedings of the Royal Society of London. Series B: Biological Sciences 266: 2471–2477. https://doi.org/10.1098/rspb.1999.0948

Deng, H.W. & Lynch, M. 1996. Change of Genetic Architecture in Response to Sex. Genetics 143: 203–212. https://doi.org/10.1093/genetics/143.1.203

Duneau, D., Altermatt, F., Ferdy, J.-B.F., Ben-Ami, F. & Ebert, D. 2020. Estimation of the propensity for sexual selection in a cyclical parthenogen. bioRxiv. https://doi.org/10.1101/2020.02.05.935148.

Duneau, D., Luijckx, P., Ruder, L.F. & Ebert, D. 2012. Sex-specific effects of a parasite evolving in a female-biased host population. BMC Biology 10: 104. https://doi.org/10.1186/1741-7007-10-104

Edgar, R.C. 2004. MUSCLE: Multiple sequence alignment with high accuracy and high throughput. Nucleic Acids Research 32: 1792–1797. https://doi.org/10.1093/nar/gkh340

Fitzpatrick, J.L., Kahrl, A.F. & Snook, R.R. 2022. SpermTree, a species-level database of sperm morphology spanning the animal tree of life. Scientific Data 9: 1–6. https://doi.org/10.1038/s41597-022-01131-w

Flössner, D. 2000. Die Haplopoda und Cladocera (ohne Bosminidae) Mitteleuropas. Backhuys Publishers, Leiden, The Netherlands.

Guindon, S., Dufayard, J.F., Lefort, V., Anisimova, M., Hordijk, W. & Gascuel, O. 2010. New algorithms and methods to estimate maximum-likelihood phylogenies: Assessing the performance of PhyML 3.0. Systematic Biology 59: 307–321. https://doi.org/10.1093/sysbio/syq010

Harmon, L.J., Weir, J.T., Brock, C.D., Glor, R.E. & Challenger, W. 2008. GEIGER: Investigating evolutionary radiations. Bioinformatics 24: 129–131. https://doi.org/10.1093/bioinformatics/btm538

Hoang, D.T., Chernomor, O., von Haeseler, A., Minh, B.Q. & Vinh, L.S. 2018. UFBoot2: Improving the ultrafast bootstrap approximation. Molecular Biology and Evolution 35: 518–522. https://doi.org/10.1093/molbev/msx281

Immler, S., Pitnick, S., Parker, G.A., Durrant, K.L., Lüpold, S., Calhim, S., et al. 2011. Resolving variation in the reproductive tradeoff between sperm size and number. Proceedings of the National Academy of Sciences of the United States of America 108: 5325–5330. https://doi.org/10.1073/pnas.1009059108

Kahrl, A.F., Snook, R.R. & Fitzpatrick, J.L. 2021. Fertilization mode drives sperm length evolution across the animal tree of life. Nature Ecology and Evolution 5: 1153–1164. https://doi.org/10.1038/s41559-021-01488-y

Kalyaanamoorthy, S., Minh, B.Q., Wong, T.K.F., Von Haeseler, A. & Jermiin, L.S. 2017. ModelFinder: Fast model selection for accurate phylogenetic estimates. Nature Methods 14: 587–589. https://doi.org/10.1038/nmeth.4285

Kishino, H. & Hasegawa, M. 1989. Evaluation of the maximum likelihood estimate of the evolutionary tree topologies from DNA sequence data, and the branching order in hominoidea. Journal of Molecular Evolution 29: 170–179. https://doi.org/10.1007/BF02100115

Kishino, H., Miyata, T. & Hasegawa, M. 1990. Maximum likelihood inference of protein phylogeny and the origin of chloroplasts. Journal of Molecular Evolution 31: 151–160. https://doi.org/10.1007/BF02109483

Lee, D., Nah, J.S., Yoon, J., Kim, W. & Rhee, K. 2019. Live observation of the oviposition process in Daphnia magna. PLoS ONE 14: 1–9. https://doi.org/10.1371/journal.pone.0224388

Lüpold, S. & Pitnick, S. 2018. Sperm form and function: What do we know about the role of sexual selection? Reproduction 155: R229–R243. https://doi.org/10.1530/REP-17-0536

Minh, B.Q., Schmidt, H.A., Chernomor, O., Schrempf, D., Woodhams, M.D., Von Haeseler, A., et al. 2020. IQ-TREE 2: New Models and Efficient Methods for Phylogenetic Inference in the Genomic Era. Molecular Biology and Evolution 37: 1530–1534. https://doi.org/10.1093/molbev/msaa015

Nguyen, L.T., Schmidt, H.A., Von Haeseler, A. & Minh, B.Q. 2015. IQ-TREE: A fast and effective stochastic algorithm for estimating maximum-likelihood phylogenies. Molecular Biology and Evolution 32: 268–274. https://doi.org/10.1093/molbev/msu300

Niksirat, H., Kouba, A. & Kozák, P. 2014. Post-mating morphological changes in the spermatozoon and spermatophore wall of the crayfish Astacus leptodactylus: Insight into a non-motile spermatozoon. Animal Reproduction Science 149: 325–334. https://doi.org/10.1016/j.anireprosci.2014.07.017

Paradis, E. & Schliep, K. 2019. Ape 5.0: An environment for modern phylogenetics and evolutionary analyses in R. Bioinformatics 35: 526–528. https://doi.org/10.1093/bioinformatics/bty633

Parker, G.A. 1982. Why are there so many tiny sperm? Sperm competition and the maintenance of two sexes. Journal of Theoretical Biology 96: 281–294. https://doi.org/10.1016/0022-5193(82)90225-9

Pennell, M.W., Eastman, J.M., Slater, G.J., Brown, J.W., Uyeda, J.C., Fitzjohn, R.G., et al. 2014. Geiger v2.0: An expanded suite of methods for fitting macroevolutionary models to phylogenetic trees. Bioinformatics 30: 2216–2218. https://doi.org/10.1093/bioinformatics/btu181

Pitnick, S., Hosken, D.J. & Birkhead, T.R. 2009. Sperm morphological diversity. In: Sperm Biology, pp. 69–149. https://doi.org/10.1016/B978-0-12-372568-4.00003-3

Popova, E. V., Petrusek, A., Kořínek, V., Mergeay, J., Bekker, E.I., Karabanov, D.P., et al. 2016. Revision of the old world Daphnia (Ctenodaphnia) similis group (Cladocera: Daphniidae). Zootaxa 4161: 1–40.

R Core Team. 2020. R: A language and environment for statistical computing. Vienna, Austria.

Ramm, S.A., Schärer, L., Ehmcke, J. & Wistuba, J. 2014. Sperm competition and the evolution of spermatogenesis. Molecular Human Reproduction 20: 1169–1179. https://doi.org/10.1093/molehr/gau070

Revell, L.J. 2012. phytools: An R package for phylogenetic comparative biology (and other things). Methods in Ecology and Evolution 3: 217–223. https://doi.org/10.1111/j.2041-210X.2011.00169.x

Roldan, E.R.S. 2019. Sperm competition and the evolution of sperm form and function in mammals. Reproduction in Domestic Animals 54: 14–21. https://doi.org/10.1111/rda.13552

Roulin, A.C., Routtu, J., Hall, M.D., Janicke, T., Colson, I., Haag, C.R., et al. 2013. Local adaptation of sex induction in a facultative sexual crustacean: insights from QTL mapping and natural populations of Daphnia magna. Molecular Ecology 22: 3567–3579. https://doi.org/10.1111/mec.12308

Seidl, M.D., Pirow, R. & Paul, R.J. 2002. Water fleas (Daphnia magna) provide a separate ventilatory mechanism for their brood. Zoology 105: 15–23. https://doi.org/10.1078/0944-2006-00050

Shimodaira, H. 2002. An approximately unbiased test of phylogenetic tree selection. Systematic Biology 51: 492–508. https://doi.org/10.1080/10635150290069913

Shimodaira, H. & Hasegawa, M. 1999. Multiple comparisons of log-likelihoods with applications to phylogenetic inference. Molecular Biology and Evolution 16: 1114–1116.

Strimmer, K. & Rambaut, A. 2002. Inferring confidence sets of possibly misspecified gene trees. Proceedings of the Royal Society B: Biological Sciences 269: 137–142. https://doi.org/10.1098/rspb.2001.1862

Wingstrand, K.G. 1978. Comparative spermatology of the Crustacea Entomostraca; 1, Subclass Branchiopoda. Biologiske Skrifter 22: 1–67.

Wuerz, M., Huebner, E. & Huebner, J. 2017. The morphology of the male reproductive system, spermatogenesis and the spermatozoon of Daphnia magna (Crustacea: Branchiopoda). Journal of Morphology 278: 1536–1550. https://doi.org/10.1002/jmor.20729

Xu, S., Ackerman, M.S., Long, H., Bright, L., Spitze, K., Ramsdell, J.S., et al. 2015. A male-specific genetic map of the microcrustacean Daphnia pulex based on single-sperm whole-genome sequencing. Genetics 201: 31–38. https://doi.org/10.1534/genetics.115.179028

Zaffagnini, F. 1987. Reproduction in Daphnia. In: Daphnia (R. H. Peters & R. De Bernadi, eds), p. 280. Istituto Italiano di Idrobiologia, Pallanza.

